# Smart bone plates can monitor fracture healing

**DOI:** 10.1101/366039

**Authors:** Monica C. Lin, Diane Hu, Meir Marmor, Safa T. Herfat, Chelsea S. Bahney, Michel M. Maharbiz

## Abstract

**One Sentence Summary:** Electrical impedance measurements using microscale sensors implanted in two mouse fracture models tracked longitudinal differences between individual mice with proper healing and mice experiencing poor healing, laying the groundwork for translation to the clinic through integration into fracture fixation implants (i.e. instrumented bone plates).

**Abstract:** There are currently no standardized methods for monitoring fracture healing. While histological studies can clearly identify the tissues found in the four stages of repair, in practice surgeons rely on X-ray, which is only useful at later stages of healing after mineralization has occurred. As electrical impedance spectroscopy (EIS) has previously been used to distinguish tissue types during healing, we hypothesized that microscale sensors embedded in the fracture callus could track the changing tissue with high sensitivity. Using *in vivo* mouse fracture models, we present the first evidence that microscale instrumented implants provide a route for post-operative fracture monitoring. In this study, we implanted sensors in mouse long bone fractures fixed with either external fixators or bone plates. EIS measurements taken across two electrodes implanted in the fracture gap were able to track longitudinal differences between individual mice with proper healing and mice experiencing poor healing. We additionally present an equivalent circuit model that combines the EIS data in order to classify healing states of fractures. Lastly, we show that EIS measures are strongly correlated with standard µCT measures of healing and that these correlations validate clinically-relevant operating frequencies for implementation of this technique. The data from these two models demonstrate that this technique can be translated to the clinic through integration into current fracture management strategies such as bone plating, providing physicians with quantitative information about the state of a fracture to guide clinical decision-making for patients.

## Introduction

Musculoskeletal injuries are among the most disabling and costly conditions in the United States, with the total number of bone fractures ranging from 12 to 15 million per year *(1)*. Treatment of these fractures represents a significant burden on the U.S. healthcare system, with hospitalization costing an estimated $23.4 billion in 2004 *(2)*. Determining when a fracture is healed is crucial to making correct clinical decisions for patients, but a survey of over 400 orthopaedic surgeons revealed that there is no consensus in defining both clinical and radiographic criteria for outcome measures *(3)*. Multiple studies cite the lack of standardized methods for assessing fracture union *(4–6)*.

Radiographic imaging and clinical evaluation are the two standard methods of monitoring bone fracture healing in the clinic. Plain X-ray radiographs are often used to assess fractures, but studies have shown that these correlate poorly with bone strength, do not define union with enough accuracy, and are unreliable for determining the stage of fracture repair. Other radiographic techniques such as computed tomography (CT), dual energy X-ray absorptiometry (DEXA), and ultrasound can offer improved diagnostic capabilities but have limited use in the clinic due to cost, high radiation dose, decreased accuracy around metal implants, or dependence on operator expertise. *(7, 8)* New monitoring tools include mechanical assessment and serologic markers, but these technologies are still under development and tend only to be used in academic research settings *(3)*. In addition, mechanical tests typically rely on a patient to be weight-bearing, limiting their utility in understanding early stages of healing *(9)*. Using serologic biomarkers as early predictors of fracture healing offers great promise, but appropriate markers correlating to the biological progression of healing have not yet been identified *(10)*. As such, physical examination by a physician is still the most relied upon technique to determine progression of fracture healing. Patients are examined for local signs of infection, ability to weight-bear, localized tenderness to palpation, and extent of pain. However, patient-reported questionnaires result in imprecise assessments, and physician evaluation is subjective and depends on experience *(3)*.

Fracture healing proceeds through a combination of two different pathways: intramembranous (direct) and endochondral (indirect) ossification *(11, 12)*. At the onset of fracture injury, formation of a hematoma around the trauma contains the fracture debris and initiates a pro-inflammatory cascade (Stage 1) *(13, 14)*. Following this, the primary healing pathway is largely determined by stability at the fracture site. In highly stable areas, cells originating from the endosteum and periosteum undergo intramembranous ossification to directly form osteoblasts (bone) *(15, 16)*. However, within the fracture gap, new bone forms indirectly through endochondral ossification, where periosteal progenitor cells undergo chondrogenic differentiation to form a cartilage intermediate (Stage 2) *(15, 17)*. Hypertrophic maturation of chondrocytes then promotes mineralization and leads to conversion into trabecular bone (Stage 3) *(18)*, and finally remodels into functional cortical bone (Stage 4). These four defined stages of healing are well characterized histologically *(12, 18, 19)*, but early stages in particular are not detectable by standard methods of monitoring like X-ray that rely on mineralization of bone.

Monitoring fracture healing is an area of active academic research, but most work has focused around obtaining mechanical feedback correlating strain measurements to bone strength *(20, 21)*. In this study, we utilize electrical techniques to characterize progression of fracture repair, building on previous work that has measured electrical changes in cells and tissue *(22–24)*. Electrically, tissue can be modelled as a combination of resistive and capacitive effects. The ion-rich intra- and extracellular matrices conduct charge and thus can be modeled as resistances, while double-layered cell membranes act as barriers to charge flow and can be modeled as capacitances or constant phase elements (CPE) *(25)*. Electrical impedance spectroscopy (EIS) measures the frequency-dependent combination of these components, describing opposition to the flow of electrical current through a material. Complex impedance *Z* can be expressed in polar form as

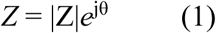
with magnitude |Z| and phase angle θ, or in Cartesian form as

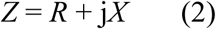
with resistance R and reactance X.

There is an extensive body of literature *(25–28)* demonstrating the use of this so-called “bioimpedance” method to quantitatively characterize cellular changes, primarily reflecting cell membrane integrity, cell volumes, and the conductivity of intra- and extracellular components *(28)*. Previous work has also detailed the use of impedance spectroscopy to describe many biological tissues *(28–31)*, including the dielectric and conductive properties of porous and dense bone *(32–34)*. We have also previously shown the feasibility of using EIS to distinguish tissue composition within the fracture callus using *ex vivo* model systems *(35, 36)*. In addition, prior work has used EIS to evaluate bone fractures in *ex vivo* and *in vivo* models *(37–42)*, but suffers from restricted detection due to noise from surrounding soft tissue, limited sensitivity due to electrode placement, and inadequate histological analysis.

Here, we present the development and testing of microscale EIS sensors designed to measure the electrical properties of the fracture callus longitudinally during healing in clinically relevant murine fracture models. To our knowledge, this is the first study to implant microscale sensors directly in the fracture gap, enabling local measurements of the changing fracture callus. We first validated the ability of our sensors to distinguish between healing and non-healing fractures in an external fixator model, where tibia defects were stabilized using a miniaturized Ilizarov external fixator. We then applied this technique to a bone plate model, where femur fractures were stabilized internally with a bone plate. Using the translationally relevant bone plate model where electrodes are embedded within fracture gaps, we find that the frequency spectra of impedance measurements are robustly correlated with quantified measures of bone volume and bone mineral density. Furthermore, this data demonstrates the use of EIS as a simple and low-cost method to provide clinically relevant information over the course of fracture healing, which can be readily integrated into clinical flows for fracture management, such as bone plates. With the global orthopaedic devices market expected to reach $41.2 billion by 2019 *(43)*, we believe there is vast potential for a smart implant and present this study as validation of EIS for integration into fracture management.

## Results

### Impedance distinguishes between fractures versus critical-sized defects in an external fixator model

As a proof-of-concept study to measure impedance of a changing fracture callus over the course of healing, we implanted small sensors into broken tibias stabilized in an external fixator model (Fig. 1A). This enabled us to quickly validate our technique using commercially-fabricated sensors protected by the large metal construct of the Ilizarov device that secured the leg. We began by implanting 250 µm diameter FR4 (a glass-reinforced epoxy laminate material) sensor pins (Fig. 1B-D) into 0.5 mm (N=6) and 2 mm (N=5) defects to determine if we could distinguish EIS trends between previously established healing versus non-healing fracture models *(44–46)*. EIS measurements were taken twice a week over a frequency range from 20 Hz to 1 MHz with mice sacrificed on days 7, 14, and 28. Mice with 0.5 mm defect sizes showed evidence of healing with a heterogeneous fracture callus composed primarily of cartilage and new trabecular bone 14 days post-injury (Fig. 2A). However, mice with 2 mm defect sizes contained only undifferentiated fibrous tissue within the fracture gap, demonstrative of poor healing at the same time point (Fig. 2B). Histology images have been false-colored to aid in interpretation of tissue composition, and original histology images are provided in Fig. S1A-B. Electrical resistance (R) over the number of days post-fracture is shown in Fig. 2C. Linear regression analyses indicate that there is a significant positive relationship between R and time in the mice with 0.5 mm defects (p < 0.0001), while there is no correlative relationship between R and time in the mice with 2 mm defects. By day 28, histology for the 0.5 mm defects show new bone formation, while 2 mm critical-sized defects result in an atrophic nonunion with complete absence of bony bridging

**Figure 1.**
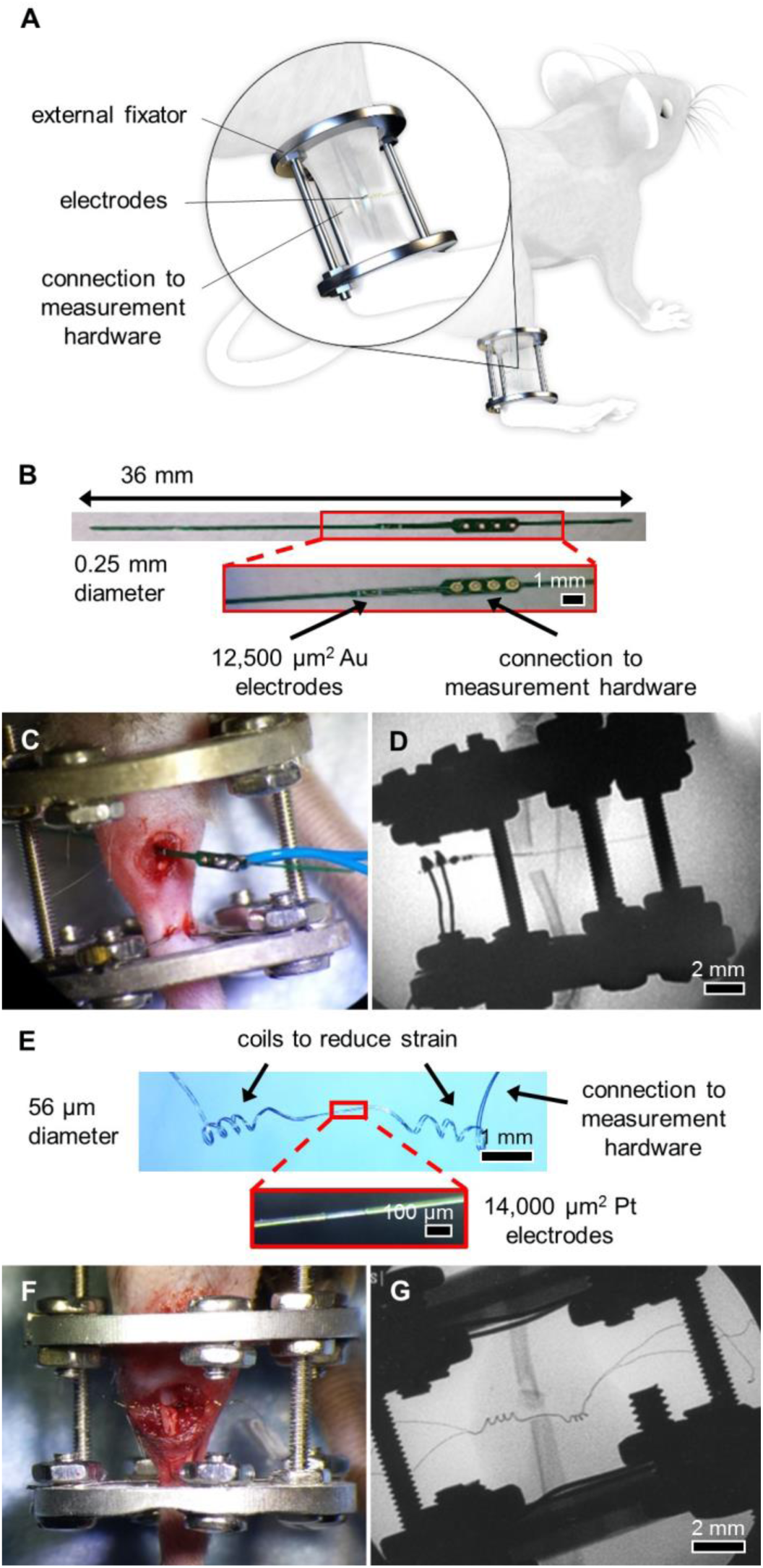
System overview and sensors for an external fixator model. **(A)** System overview of sensor embedded in mouse tibia fracture, where the injury is stabilized with an external fixator. **(B)** 0.25 mm diameter sensor fabricated on an FR4 substrate, with electroless-nickel immersion gold (ENIG) surface electrodes and large vias outside the leg to connect to measurement hardware. Sensors were implanted in externally-fixed mice tibias with 0.5 mm and 2 mm defects. **(C)** Photograph of open surgery performed to implant 0.25 mm sensor in the external fixator model. Surgical site was closed following sensor placement. **(D)** Fluoroscopy image of implanted 0.25 mm sensor in a 2 mm defect model. **(E)** 56 µm diameter sensor assembled using platinum wire, with recording sites exposed by a CO_2_ laser and coils added to provide strain relief. **(F)** Photograph of open surgery performed to implant 56 µm sensor in the external fixator model. Surgical site was closed following sensor placement. **(G)** Fluoroscopy image of implanted 56 µm sensor in a 0.5 mm defect model.

**Figure 2.**
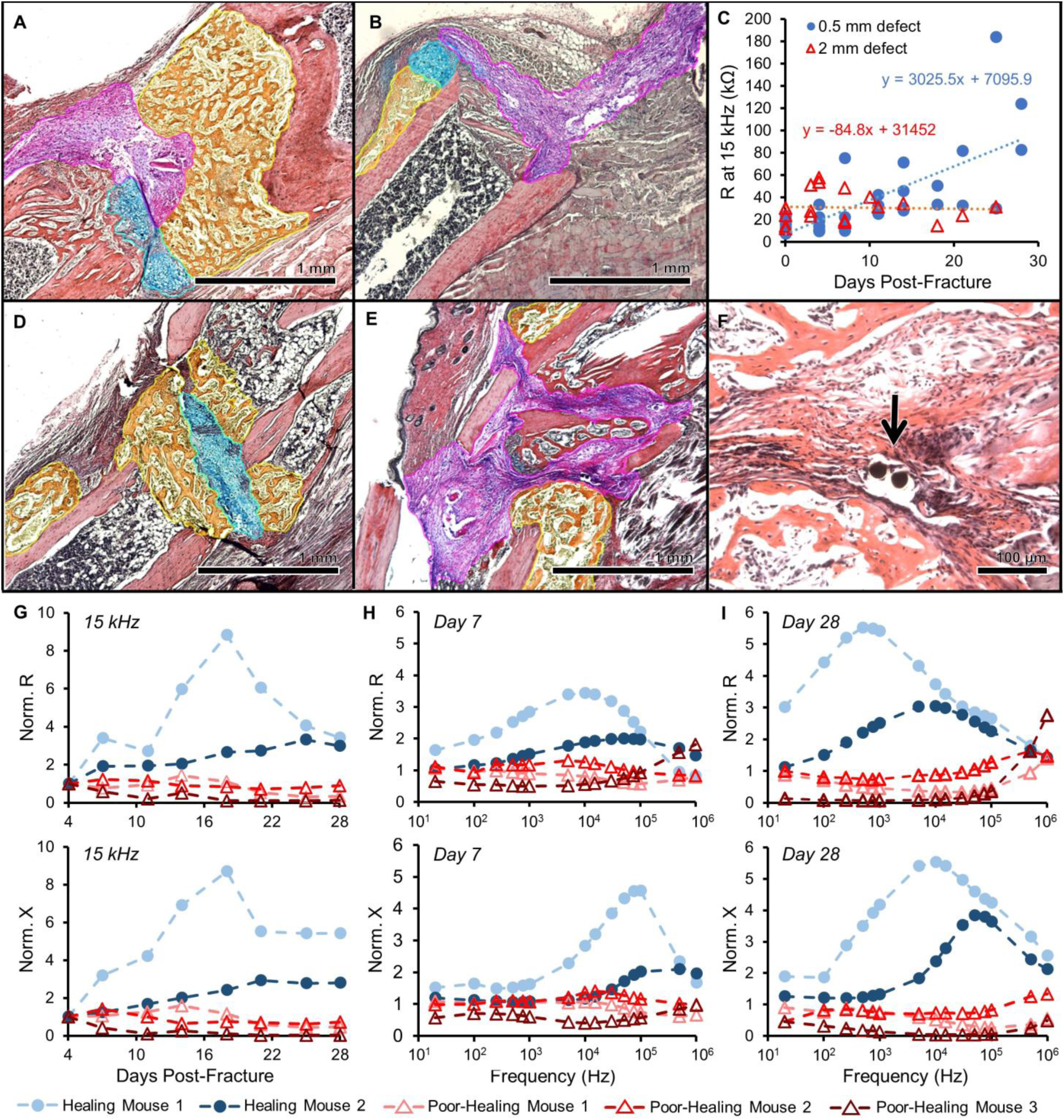
Impedance data distinguishes healing and poorly healing tibia fractures in an external fixator model. Histology sections in this image are stained with Hall’s and Brunt’s Quadruple (HBQ) stain and false-colored to aid interpretation of tissue composition. Blue = cartilage, yellow = trabecular bone, and purple = fibrous/amorphous tissue. Original red color = cortical bone, black/white area = bone marrow. **(A)** Representative histology section for an externally-fixed 0.5 mm defect at 14 days post-fracture; the fracture gap is clearly bridged by cartilage and new trabecular bone. **(B)** Representative histology section for an externally-fixed 2 mm critical-sized defect at 14 days post-fracture; the fracture gap is dominated by fibrous tissue. **(C)** Electrical resistance (R) at 15 kHz measured with a 250 µm sensor is plotted over days post-fracture for measurements taken in mice with 0.5 mm (N=6) and 2 mm (N=5) defects. Linear regression analyses determined that there is a significant positive relationship in mice with 0.5 mm defects (p < 0.0001), while there is no correlative relationship in mice with 2 mm defects. **(D)** Representative histology section for a healing mouse at 28 days post-fracture; the fracture gap is clearly bridged by cartilage and new trabecular bone. **(E)** Representative histology section for a poor-healing mouse at 28 days post-fracture; the fracture gap contains an overabundance of fibrous tissue. **(F)** Black arrow points to 56 µm sensors fully embedded in fracture tissue. **(G)** Electrical resistance (R) and reactance (X) normalized as a ratio to day 4 plotted over the course of fracture healing at 15 kHz. Normalized R and X both rise steadily over healing time in the healing mice, with stagnant values observed in the poor-healing mice. **(H)** Normalized R and X as a ratio to day 4 plotted over a range of frequencies at day 7 post-fracture. **(I)** Normalized R and X as a ratio to day 4 plotted over a range of frequencies at day 28 post-fracture. Marked shifts in frequency response from day 7 were observed in the healing mice, with limited change occurring in the poor-healing mice.

### Microscale sensors track changes in frequency response for healing and non-healing fractures in an external fixator model

One of the limitations of the sensor utilized for the initial validation study was that we consistently noted fibrous tissue surrounding the sensor even out to 28 days post-injury. We hypothesized that this may be due to the large size of the sensor relative to the tibia defect. Consequently, we designed and built significantly smaller electrodes (56 µm diameter platinum) for implantation, measuring roughly an order of magnitude thinner than the original FR4 sensors *(47)*.

Using these 56 µm diameter platinum (Pt) wire electrodes (Fig. 1E) implanted within fractures stabilized in the same external fixator model (Fig. 1F-G) with 0.5 mm defects, we again measured EIS from 20 Hz to 1 MHz twice per week out to 28 days post-injury in five mice. In two of the mice, histology showed clear signs of healing, with a robust callus present between the two bone ends (Fig. 2D). However, the other three mice displayed poor signs of healing, with their fracture gaps dominated by an overabundance of fibrous tissue (Fig 2E, original histology in Fig. S1C-D). To analyze EIS data as it correlated to histological evidence of healing, electrical resistance (R) and reactance (X) were normalized as a ratio to the first time point after surgery (day 4). This was done to account for the initial perturbation of surgery, variation between sensors and samples, and to enable comparison between multiple mice. Based on all EIS plots, we found the largest spread between measurement days at 15 kHz, indicating that this is the most functional frequency for detecting differences in fracture healing (Fig. 2G). This falls within the frequency range that can be identified as the beta dispersion, which is associated with the polarization of cell membranes, proteins, and other macromolecules *(28)*. At this frequency, normalized R increases with progression of fracture repair in the healing mice, but stalls in the poor-healing mice as expected. Normalized X rises above 1 over time in healing mice, and below 1 over time in poor-healing mice. Fig. 2H-I illustrates how normalized R and X as functions of frequency differ for healing and poor-healing mice from an early time point (day 7) to a late time point (day 28). The healing mice exhibit clear changes in their frequency response in both parameters relative to day 4, especially in the frequency range of 10^3^ to 10^5^ Hz. Comparatively, non-healing mice show relatively flat frequency responses relative to their initial day 4 behavior. The influence of callus formation on the frequency response of impedance measurements enables the clear differentiation between well-healing and poor-healing mice.

In the poor-healing mice, a large amount of fibrous tissue was noted and we hypothesized this was due to excess movement of the sensor relative to the fractured bone ends. By comparing fluoroscopy images taken immediately after surgery and on day 28 before euthanasia, we can determine whether or not the sensor shifted over time. In the two healing mice, day 28 X-rays showed the sensor remained unchanged and in the same location as on day 0. However, in the three poor-healing mice, the coils flanking the exposed electrodes were no longer present on the day 28 X-rays, a clear indication that the sensor moved from its original position. Yet this thin and flexible nature of the sensors also allowed them to be preserved throughout the histology process, enabling us to visualize and confirm the specific tissue surrounding the sensor (Fig. 2F).

### Impedance discriminates range of healing found in bone plate model fractures

To extend the utility of our monitoring technique to a more clinically relevant fixation procedure, we designed and built sensors for integration with a bone plate model. Bone plates are commonly used for patients with traumatic fractures that require surgical intervention with internal fixation and provide a platform from which to integrate much shorter sensors that will thus experience significantly less movement compared to the external fixator model. We designed small sensors using a flexible polyimide sensor board that could be affixed to the proximal half of commercially available murine bone plates and 25 µm-diameter platinum wires attached at the center to serve as electrodes within the fracture gap (Fig. 3A-B). A long, serpentine cable extended from the proximal end of the plate and was tunneled subcutaneously to the shoulder blades, leading to a set of connection pads that interfaced with our measurement instrumentation. To test our “smart” bone plate, an open surgery was performed to affix the plate to the mouse femur and the sensor wire was inserted into a reproducible (<0.25 mm) fracture gap (Fig. 3C-D). Surgery was also performed on control mice using the same procedure but without fracturing the bone; in this case, the sensor was placed beside the bone and muscle was sutured over it. 2-point impedance measurements were taken from 20 Hz to 1 MHz between two 700 µm^2^ electrodes three times per week (Fig. S2), with mice euthanized at 12 (N=3 fracture) or 26 (N=5 fracture, N=2 control) days post-surgery for histology and micro-Computed Tomography (µCT) analysis.

**Figure 3.**
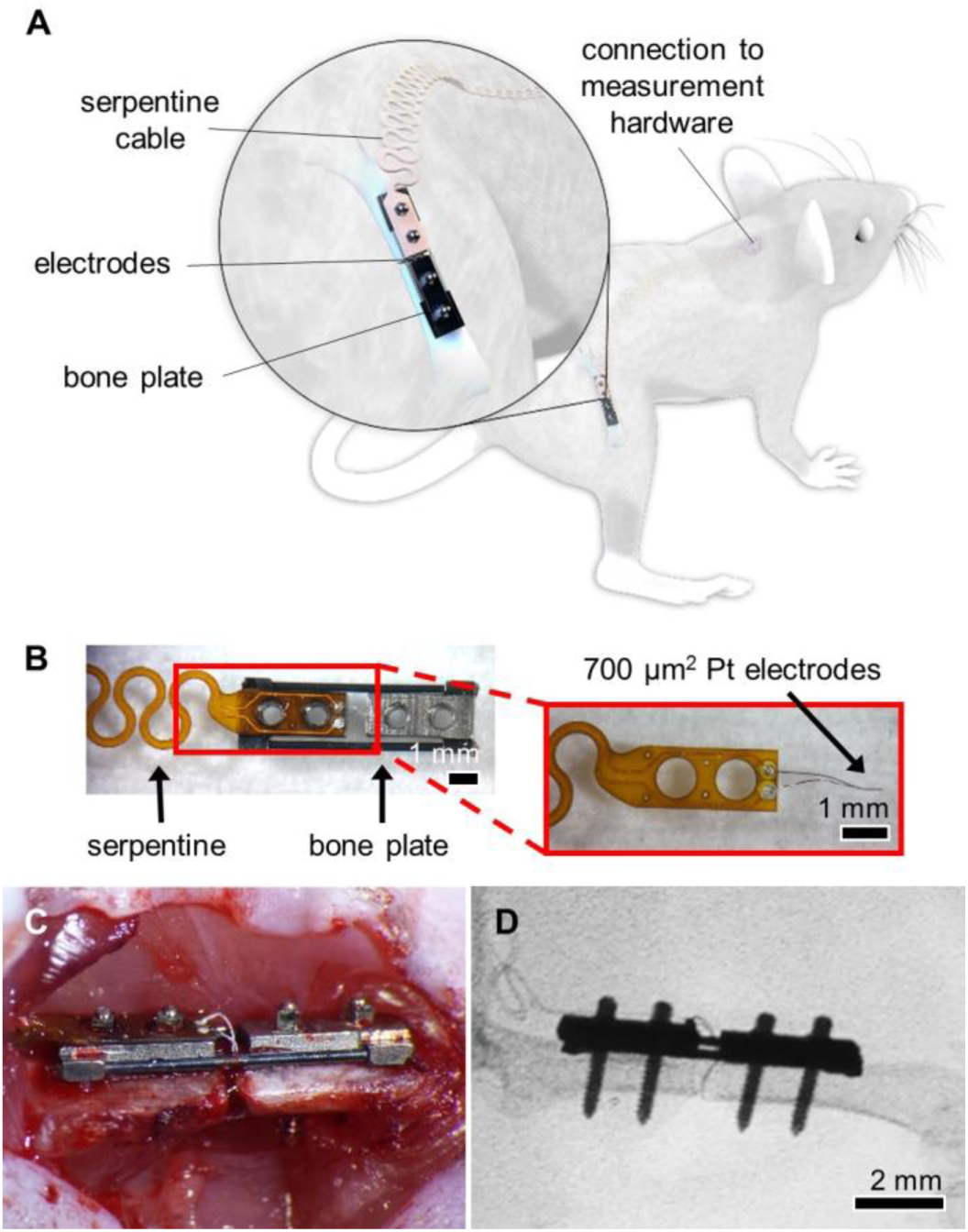
System overview and sensor for a bone plate model. **(A)** System overview of sensor embedded in mouse femur fracture, where the injury is stabilized with a bone plate. **(B)** Sensors fabricated on a polyimide substrate with 700 µm2 platinum (Pt) electrodes spaced 0.5 mm apart. Sensors were affixed to the proximal half of the bone plate, with a long flexible cable extending off the proximal end. The serpentine pattern repeated for the length of the cable, ending in two vias that served as connectors that link to the measurement hardware. **(C)** Photograph of open surgery performed to implant a bone plate and affixed sensor to stabilize a femur fracture. Surgical site was closed following sensor placement. **(D)** Fluoroscopy image of implanted Pt sensor in the fracture gap.

The mice experienced varied degrees of healing over the course of the study. Some fractures healed very well, with complete bony bridging seen by histology (Fig. 4A), X-ray (Fig. 4C), and µCT (Fig. 4D). While all samples showed evidence of a healing response, in some cases a bony callus did not bridge the fractured bone ends (Fig. 4E); this could also be seen to some extent in X-ray (Fig. 4G) and µCT (Fig. 4H). Original histology images are provided in Fig. S3. We were again able to define the exact tissues immediately surrounding the sensor in each sample, regardless of the global callus composition. In Fig. 4B, we see that the sensor is completely embedded in new bone, while in Fig. 4F, the sensor is at the boundary between new bone and fibrous tissue. Representative histology images for each sample are shown in Fig. 5 to indicate resultant healing state (original images in Fig. S4), and the complete set of X-ray and µCT images are provided in Fig. S5. In the control mice, the sensor stayed between the unfractured bone and surrounding muscle tissue, but there was evidence of fibrous tissue surrounding it (Fig. 5I, Fig. S6).

**Figure 4.**
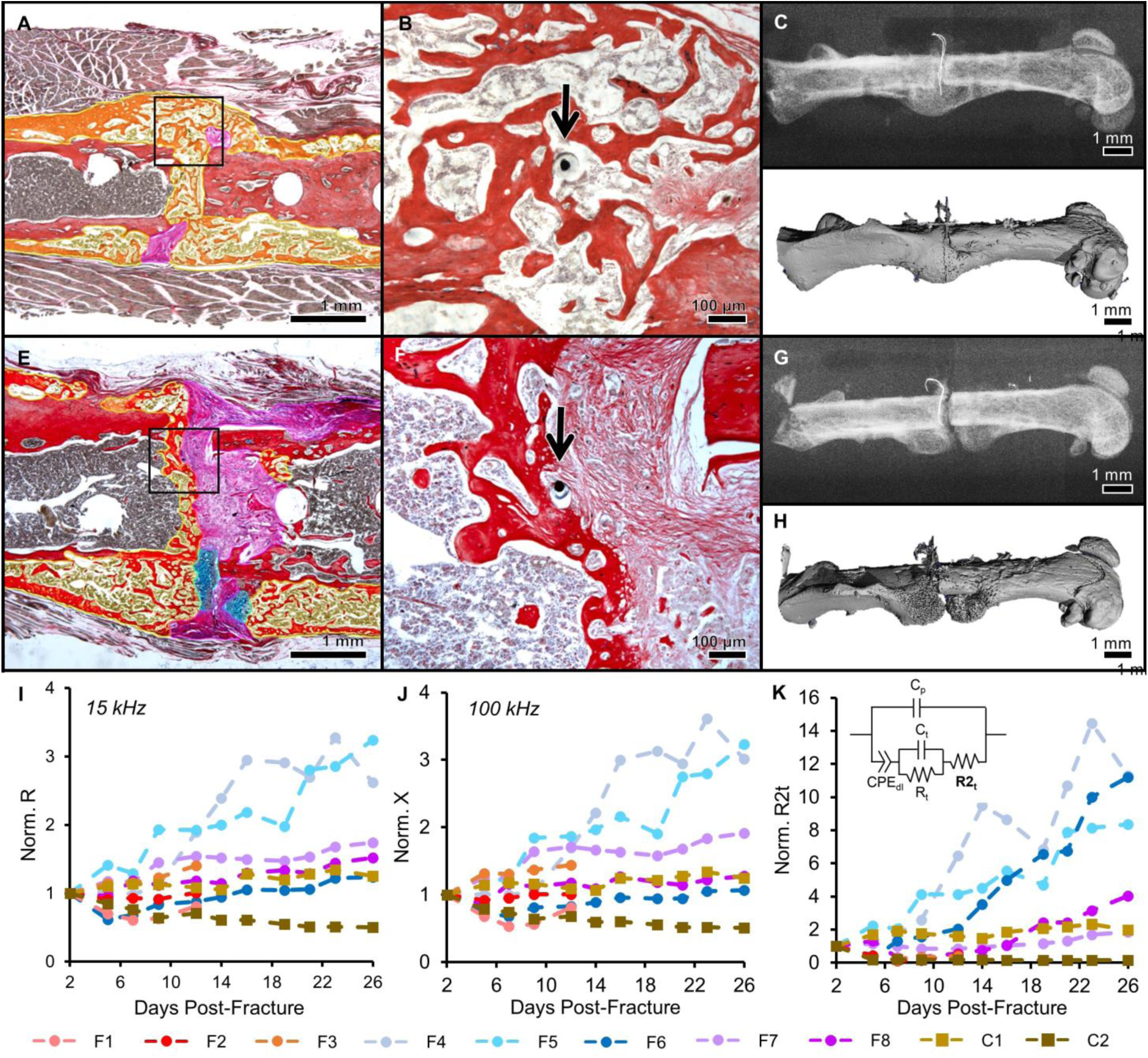
Impedance data distinguished femur fractures completely healed from those with varied healing in a bone plate model. Histology sections are stained and false-colored like in Fig. 2. **(A)** Representative histology section of a well-healed mouse at day 26; a large bony callus has formed that completely bridges the two fracture ends. Black box outlines position of the high-magnification image in (B). **(B)** High magnification image of (A) with black arrow pointing to the electrode fully-integrated in new trabecular bone. **(C)** X-ray radiograph of well-healed fracture on day 26. **(D)** Surface rendered, three-dimensional µCT image of well-healed fracture on day 26. **(E)** Representative histology section of a mouse with mixed healing at day 26; the fracture callus includes cartilage, fibrous tissue, as well as trabecular bone. Black box outlines position of the high-magnification image in (F). **(F)** High magnification image of (E) with black arrow pointing to the electrode fully-embedded in the callus. It is surrounded by a mixture of new trabecular bone and fibrous tissue. **(G)** X-ray radiograph of fracture with mixed healing on day 26. **(H)** Surface rendered, three-dimensional µCT image of fracture with mixed healing on day 26. **(I)** R (normalized as a ratio to day 2) at 15 kHz plotted as a function of days post-fracture. Data markers and lines are colored according to degree of healing – shades of red for mice sacrificed at day 12 (F1, F2, and F3), shades of blue for mice sacrificed at day 26 that healed well (F4, F5, and F6), shades of purple for mice sacrificed at day 26 that healed poorly (F7 and F8), and shades of brown for control mice sacrificed at day 26 (C1 and C2). Normalized R clearly rises at a faster rate in the two mice with complete bony calluses, F4 and F5. **(J)** X (normalized as a ratio to day 2) at 100 kHz plotted as a function of days post-fracture. Normalized X clearly rises at a faster rate in the two mice with complete bony calluses, F4 and F5. **(K)** Impedance data at all measured frequencies is fit to an equivalent circuit model (inset), and the R2t parameter is extracted, normalized as a ratio to day 2, and plotted as a function of days post-fracture. This analysis is able to clearly distinguish the samples that are classified as union by orthopedic surgeons (F4, F5, and F6 in Table 1).

**Figure 5.**
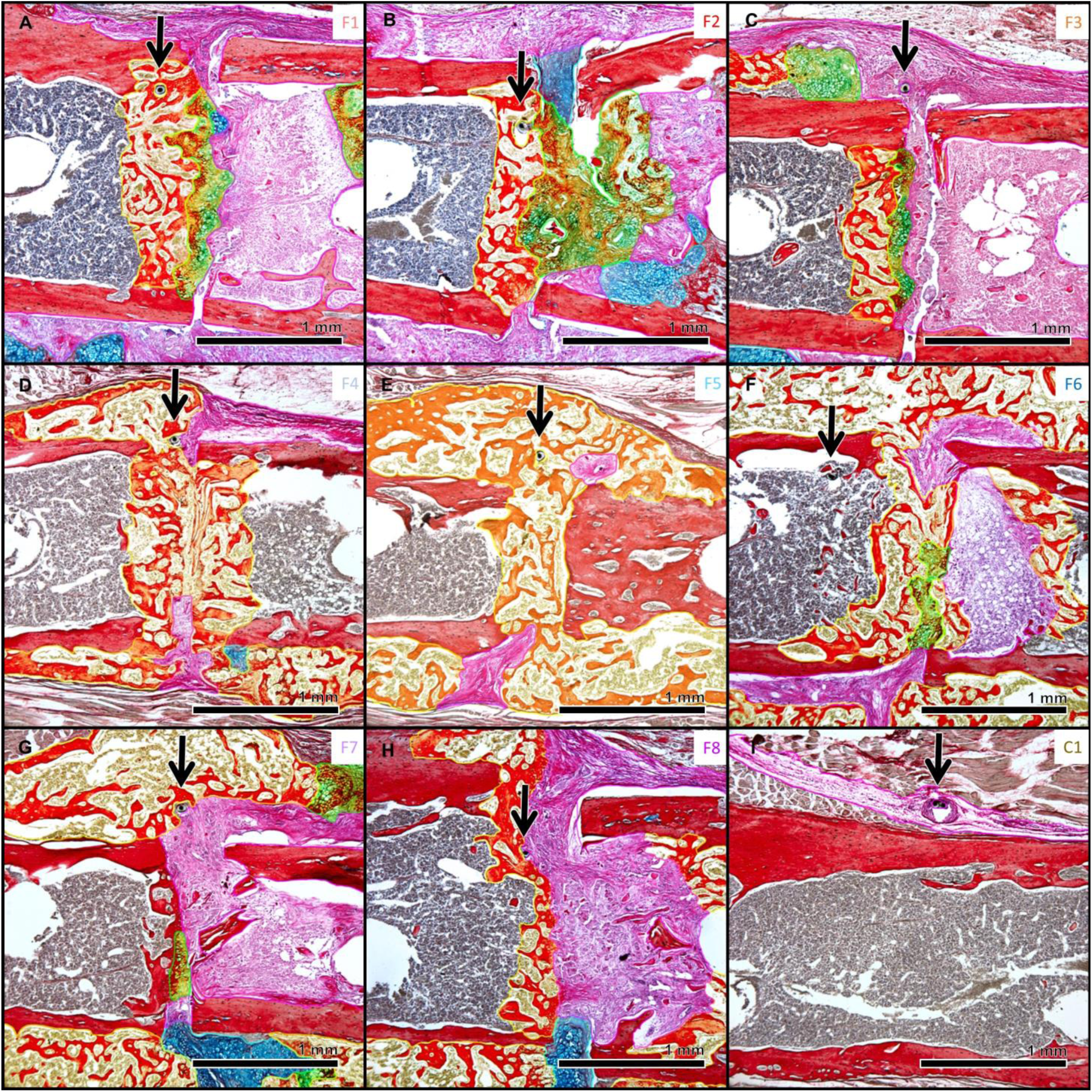
Histology sections for all bone plate model mice. Histology sections are stained and false-colored like in Fig. 2. Green = cartilage converting to bone. Black arrows point to sensors embedded in the fracture callus. Sample labels (upper right corner) are colored to match the graphs in Fig. 4I-K. **(A-C)** Histology sections for mice sacrificed at day 12. **(D-F)** Histology sections for mice sacrificed at day 26 with fracture calluses composed almost entirely of new trabecular bone. **(G-H)** Histology sections of mice sacrificed at day 26 that experienced mixed healing. **(I)** Representative histology section of a control mouse, where the sensors are embedded in a small amount of fibrous tissue.

The electrical resistance (R) at 15 kHz is plotted over time in Fig. 4I, and the electrical reactance (X) at 100 kHz is plotted over time in Fig. 4J, normalized as a ratio to day 2. These frequencies were found to reflect significant biological information, as discussed in later analysis. Data markers and lines were colored according to degree of healing – shades of red for mice sacrificed at day 12 (F1, F2, and F3), shades of blue for mice sacrificed at day 26 that healed well (F4, F5, and F6), shades of purple for mice sacrificed at day 26 that healed poorly (F7 and F8), and shades of brown for control mice sacrificed at day 26 (C1 and C2). Samples like F4 and F5 that clearly healed with their sensors well-embedded in new bone exhibited R and X values that grew steadily over time. Conversely, samples like F7 and F8 that had a mix of new bone and fibrous tissue in the fracture gap exhibited R and X values that did not rise as high, indicating stagnant healing. Measurements in the control samples remained relatively flat over time with small deviations due to the minor fibrotic response, as expected. By examining the normalized R and X values at specific frequencies over healing time, we can begin to discriminate between clearly healed samples and those with mixed healing.

### A model-driven fit of data provides robust classification of healing trends over time

In order to determine how the data best tracked healing, we fit the resistance and reactance data at all measured frequencies to an equivalent circuit model of biological tissue. This enabled us to extract the dominant electrical properties that explain the differences in healing that we observe. The equivalent circuit models the double-layer impedance at the electrode-electrolyte surface as a constant phase element (CPE_dl_) in series with the tissue component, and places a capacitor (C_p_) in parallel to reflect the parasitic capacitance due to coupling along the two long wire traces leading from the electrodes to the connector. The tissue component is comprised of a resistor (R_t_) and capacitor (C_t_) in parallel, in series with another resistor (R2_t_) (Fig. 4K). R_t_ represents the ionic intracellular environment, R2_t_ represents the ionic extracellular environment, and C_t_ represents the double-layer cell membranes. In our model, we held C_p_ constant at 25 µF, extracted from high-frequency measurements taken with our sensors immersed in Phosphate Buffered Saline (PBS) prior to implantation. The impedance CPE_dl_ is calculated as

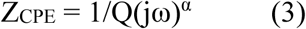
where Q is the numerical value of admittance (1/|Z|) at ω = 1 rad/s, ω is the angular frequency, and α is a constant from 0 to 1 (1 representing a perfect capacitor). We held α constant at 0.5, based on low-frequency measurements of our sensors in PBS. From a study characterizing an osseous implant using EIS *(48)*, we hypothesized that the changing impedance in our study would be most reflected in R2_t_. We thus fit R_t_ and C_t_ at an initial time point (day 2) and constrained it at that value over time. We used a custom MATLAB script to fit our data at each time point to this equivalent circuit model, interpolating 10 steps in between each time point to guide the mathematical solution and avoid jumping to non-physical values. Goodness of fits graphs are provided in Fig. S7. By normalizing R2_t_, shown in Fig. 4K, we find that this parameter accurately recapitulates healing state. The best healed samples (F4, F5, and F6) rise clearly above the samples that experienced more varied healing.

### Impedance significantly correlates with bone mineral density and percentage of new bone in a fracture callus

To understand how clinicians would characterize healing using standard classification techniques, we had five orthopaedic surgeons score the final X-rays for each sample using the modified Radiographic Union Scale for Tibial (RUST) fractures *(49)*. Each cortex is assigned a value from 1 to 4: 1 if there is no callus present, 2 if there is callus present, 3 for bridging callus, and 4 for remodeled bone where the fracture line is no longer visible. Our data excludes the anterior cortex as the sensor typically occludes the full view on an X-ray, so the total possible score ranges from 3 to 12. Each X-ray was also categorized as a union (U), nonunion (NU), or suspected nonunion (SNU); the summarized data is presented in Table 1 (raw data available in Table S1). In clearly healed cases like F4, F5, and F6, there is strong agreement between surgeons in total RUST score as evidenced by low standard deviations, and all surgeons categorized these three fractures as fully united. This corroborates our impedance data, as the R2_t_ parameter extracted from the model-driven fit also classified these three samples as well-healed. However, surgeon evaluation varied for ambiguous cases, with a wider spread in total RUST score as well as disagreement in how to categorize a fracture, substantiating the lack of a standard for assessing fracture healing by X-ray.

**Table 1.**
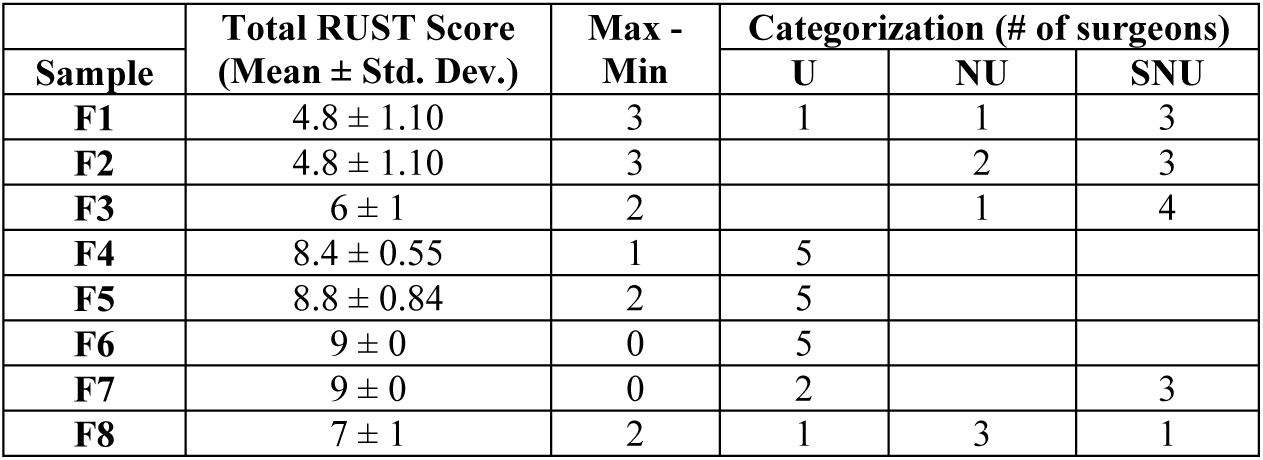
Clinician-rated RUST scores for fracture X-rays. Total RUST scores for each sample were averaged amongst five orthopaedic surgeons and presented as mean ± standard deviation. The maximum score subtracted by the minimum score for each sample was calculated to show the largest difference between surgeon evaluations. The number of surgeons who categorized each sample as union (U), nonunion (NU), or suspected nonunion (SNU) are also provided. In clearly healed cases (F4,F5,F6) there is strong agreement between clinicians, but results vary for ambiguous cases.

| Sample | Total RUST Score<br>(Mean ± Std. Dev.) | Max -<br>Min | Categorization (# of surgeons) | | |
| --- | --- | --- | --- | --- | --- |
|  |  |  | U | NU | SNU |
| F1 | 4.8 ± 1.10 | 3 | 1 | 1 | 3 |
| F2 | 4.8 ± 1.10 | 3 | | 2 | 3 |
| F3 | 6 ± 1 | 2 | | 1 | 4 |
| F4 | 8.4 ± 0.55 | 1 | 5 | | |
| F5 | 8.8 ± 0.84 | 2 | 5 | | |
| F6 | 9 ± 0 | 0 | 5 | | |
| F7 | 9 ± 0 | 0 | 2 | | 3 |
| F8 | 7 ± 1 | 2 | 1 | 3 | 1 |

In the clinic, computed tomography (CT) scans can be requested to confirm or reject suspected delayed or nonunion cases. While CT sees limited use in the clinic due to its high costs, it does enable quantitative measures that reflect biological and structural measures of healing. We quantified a number of microstructural indices from micro-computed tomography (µCT) scans of our samples to determine their degree of correlation with EIS. Preexisting bone was excluded in our analysis to only examine mineralized tissue within the fracture callus, and bone volume (BV), total volume (TV), bone mineral density (BMD), trabecular number, trabecular thickness, and trabecular separation were quantified (Table S2). Following linear regression analyses, we find that normalized resistance (R) significantly correlates with the ratio of bone volume to total volume (BV/TV), BMD, trabecular number, and trabecular thickness (Fig. 6A-D). These correlations peak at 15 kHz, with normalized R rising with increasing BV/TV (R^2^ = 0.7555, p < 0.01) and BMD (R^2^ = 0.7925, p < 0.01). In addition, normalized R at 15 kHz also has a positive relationship with trabecular number (R^2^ = 0.5641, p = 0.03) and trabecular thickness (R^2^ = 0.6417, p = 0.02), but has no correlation with trabecular separation (R^2^ = 0.3599, p = 0.12). Similar relationships are found by comparing normalized reactance (X) to these µCT indices.

**Figure 6.**
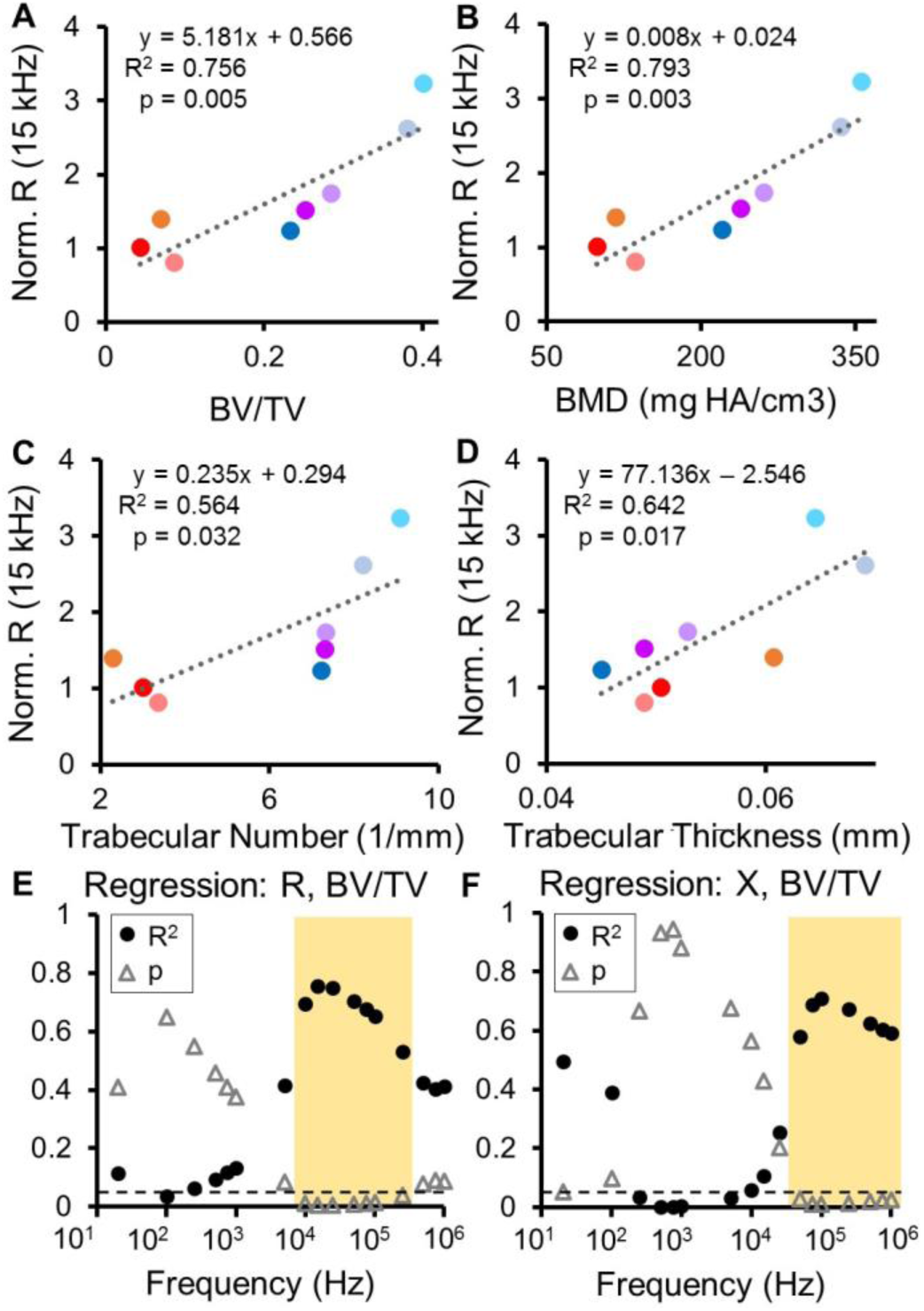
Regression analyses comparing normalized impedance data to µCT indices. **(A-D)** Normalized R at 15 kHz is significantly correlated to the ratio of bone volume to total volume (BV/TV), bone mineral density (BMD), trabecular number, and trabecular thickness. Individual markers are colored to match the corresponding samples in Fig. 4I-K. **(E-F)** Resultant R2 and p-values from regression analyses between normalized R and BV/TV as well as normalized X and BV/TV plotted as a function of frequency. Significance is set at p < 0.05 (below dashed line). Significant relationships between R and BV/TV occur from 10 kHz to 250 kHz, and significant relationships between X and BV/TV occur from 50 kHz to 1 MHz.

Importantly, by plotting the associated R^2^ and p values at every measured frequency for each pair of variables, we can pinpoint the specific frequency ranges in which impedance provides valuable information about healing. As seen in Fig. 6E, normalized R is significantly correlated to BV/TV from 10 kHz to 250 kHz (R^2^ > 0.5, p ≤ 0.04), with a peak at 15 kHz. In Fig. 6F, we see that normalized X is significantly correlated to BV/TV at a higher set of frequencies – 50 kHz to 1 MHz (R^2^ > 0.55, p < 0.03), with a peak at 100 kHz. Analogous graphs for the other µCT indices are shown in Fig. S8. Normalized R is significantly correlated with BMD from 10 kHz to 250 kHz (R^2^ > 0.55, p < 0.03), with trabecular number from 15 kHz to 100 kHz (R^2^ ≥ 0.5, p < 0.05), and with trabecular thickness from 5 kHz to 100 kHz (R^2^ > 0.45, p < 0.05). Normalized X is significantly correlated with BMD from 50 kHz to 100 kHz (R^2^ ≥ 0.6, p ≤ 0.02), and with trabecular thickness at from 25 kHz to 100 kHz (R^2^ > 0.55, p ≤ 0.03). These results are also summarized in Table S3. This regression analyses not only determines that EIS correlates to clinically-relevant measures of bone healing that would otherwise require expensive µCT evaluation, but also pinpoints the specific frequency ranges at which this is most significant.

## Discussion

Our study aimed to validate the use of EIS to develop smart fracture fixation instrumentation, with the goal of arming physicians with quantitative information about the state of fracture repair in patients. While previous research has been done utilizing impedance to evaluate bone fractures, other groups have used surface electrodes on the outside of the leg or pins drilled in the proximal and distal ends of the bone. As a result, EIS measurements in these studies reflect a global assessment with low specificity to the changing fracture callus, as signal must also travel through the muscle and skin surrounding the injured bone. To our knowledge, our study is the first to design and fabricate microscale sensors that can be implanted directly in the fracture gap intra-operatively. This enabled longitudinal EIS measurements to be taken specifically in the calli of individual mice with fractures stabilized using either external fixators or bone plates *in vivo*.

In this paper, we demonstrate the utility of EIS to track differences between fractures healing well and those healing poorly in two clinically-relevant models, laying the foundation for use of this technique to detect fracture nonunion or other complications. Beginning with an external fixator model that utilized two defect sizes, we studied the ability of EIS to distinguish between two substantially different healing pathways. The 2 mm osteotomy resulted in a critical-sized defect with no substantial indications of healing, as the fracture gap was dominated by amorphous granulation and fibrous tissue. Electrical resistance (R) did not change over time in these mice because the electrodes were consistently exposed to this amorphous mixture of fibrous tissue. In contrast, the 0.5 mm distraction resulted in an increase in R as the fracture gap included cartilage and new bone over the course of 28 days. While these results show a clear difference between impedance measurements in fractures with varied healing responses due to different defect sizes, this study was limited in that the sensor itself appeared to generate a fibrotic response.

By using a smaller, flexible sensor assembled using thin platinum wire, we were able to track EIS throughout the time course of fracture healing with decreased disruption to the healing process in both an external fixator model and bone plate model. From these two surgical models, a range of healing responses was produced from atrophic nonunion to full bridging. To understand the correlation of EIS measurements with healing, we also assessed these fractures with the full range of standard clinical and laboratory techniques, including histology, X-ray with assigned RUST scores, and µCT. Due to the small size of our sensors, histology could confirm not only the tissue composition across the entire fracture gap, but also identify the exact tissue the sensor was embedded in. While motion of the sensor in some samples likely contributed to atrophic nonunion, histological evidence showed that our microscale sensors were well integrated within the fracture tissue and when appropriately immobilized, did not prevent complete bony bridging between fractured bone ends. Full integration of our sensors within fracture calli was critical for taking local measurements of healing tissue over time in individual mice without disturbing the repair process.

We found that impedance between mice that healed well and those that healed poorly deviated quickly, with electrical resistance (R) and reactance (X) rising at a faster rate in well-healed fractures. This indicates that impedance may be able to rapidly identify stalled healing as compared to a traditional technique such as X-ray, which is only able to detect mineralized tissue at late healing stages. Resistance reflects the ability of a material to conduct charge – a highly conductive material will exhibit a small resistance, while a less conductive material will have a larger resistance. As the dominant tissue in a fracture callus transitions from highly conductive blood and cartilage to less conductive trabecular and cortical bone, our sensors confirm that R rises rapidly over time in mice that exhibited a clear healing response. Reactance describes effects that store energy such as charge separation across cell membranes (which act as barriers to charge flow and look capacitive). Highly cellular tissues have a higher capacity for energy storage and thus a larger X, particularly at higher frequencies. Because bone is comprised of multiple matrix layers that can be modeled as capacitors and marrow is highly cellular, we see a sharp rise in X in the mice with bridging calli, in contrast to more moderate changes in mice with poor healing. Both R and X stayed roughly flat in the control samples, in which no fracture was made, with minor deviations over time due to a small fibrous tissue response encapsulating the sensor. We also see that the greatest spread in R and X across days occurs in the 1 kHz to 100 kHz frequency range, and formation of a robust callus influences the frequency response of the measurement. This reveals the importance of spectroscopic measurements, which enable us to precisely track where changes arise in the frequency range over healing time. These results align with previous work in cadaveric and *ex vivo* mouse models *(35, 36)* that describe resistive components increasing over fracture healing time (blood to cartilage to bone), and reactive components rising as cartilage converts to bone. This assessment is corroborated by EIS data collected in both the external fixator and bone plate models, signifying the consistency of this technique in two different fracture fixation schemes.

By fitting our data to an equivalent circuit model, we were able to combine the effects of resistance and reactance changes at all measured frequencies and determine robust measures of trends over time. By simplifying the complex physical components and processes present in healing fractures to electrical components that suitably approximate tissue *(26, 48)*, our analysis found that the R2_t_ term describing the extracellular environment was able to clearly separate the mice with bony bridging from those without. One of the mice, F6, developed an extremely large callus with a lot of new trabecular bone as well as a pocket of amorphous tissue. This is a unique case in which the fracture was clinically considered healed (Table 1), but still had unexpected fibrous tissue within the gap. While the R and X values for this sample were grouped with mice that had varied healing, fitting to our equivalent circuit model was able to pull out all the fractures that would be clinically categorized as unions. Although the model is an approximation of the biology occurring between the two electrodes in the fracture gap, it allowed us to obtain robust measures of change which employed all of the impedance data in a combined fashion to help explain the differences in healing that we observed. With validation from a larger clinical trial, this work could be used as a platform to create prognostic tools in the clinic to predict and differentiate proper healing and delayed or nonunion.

In mice with clearly healed fractures, all techniques showed strong agreement in classifying the fractures as united. However, in cases with a more heterogeneous healing response, there was no consensus among the five surgeons, with up to 3 points difference in the total RUST score. Notably, there were two cases that were categorized as “union”, “nonunion”, and “suspected nonunion” depending on the surgeon being asked. This further emphasizes the need for an objective method to quantify the degree of repair in a healing fracture, as interpretation of X-ray can vary amongst surgeons, potentially due to factors like medical training, experience, etc. In order to correlate EIS data to more accurate measures of healing, we performed µCT. Additional quantitative data derived from analyzing µCT revealed the sensitivity of EIS measurements to relevant parameters of healing, including BV/TV, BMD, trabecular number, and trabecular thickness. Our regression analyses show that both normalized R and X have strong positive relationships with these four µCT indices within specific frequency ranges. These correlations peak in significance at 15 kHz for R and at 100 kHz for X, reflecting the primary frequency of interest to understand mineralization of the fracture callus. As we’ve described, R and X rise as cartilage is converted to bone within a fracture callus. As this healing progresses, the total amount of mineralized bone as a ratio of the total callus volume will also rise. Furthermore, bone mineral density rises correspondingly as chondrocytes become mineralized and then convert to bone. Likewise, as fractures transition from a solid cartilage callus to trabecular bone, we expect trabecular number and thickness to increase. Our impedance measurements are able to track these changes with a high degree of sensitivity, providing valuable information that would otherwise require the extra expense and radiation exposure associated with a CT scan.

While mice provided a well-established model of fracture healing that closely mimics the same process in humans, one of the biggest challenges was designing small, wired implants within tight size constraints. Since the tibia and femur bones in a mouse measure approximately 1-1.5 mm in diameter, this study implanted only the electrodes at the fracture site, with additional instrumentation to take impedance measurements located outside the animal. The flexibility of metal wires at this small size (25 to 60 µm diameter) permitted movement of some sensors, which we hypothesize occurred due to soft tissue movement near the fracture or callus formation effectively pushing the sensor. While we observed negligible changes in inter-electrode distance, future designs in a larger model should aim to eliminate potential fluctuations, possibly by using stiffer electrodes. In addition, while smaller sensors improved integration in the fracture tissue, they had higher impedance and thus possibly reduced the signal-to-noise ratio (SNR) of our measurements. Future designs could increase electrode surface area while restricting overall size to limit this effect. Translating this technology to patients will involve development of electronics to integrate wireless data transfer, which would also enable more frequent measurements. For patients already undergoing surgery to stabilize a fracture with instrumentation, a sensor could be added intra-operatively as a simple addition to standard surgical hardware to provide the patient and physician with quantitative information on the state of fracture healing. This would provide detailed information to physicians and patients about the healing trends over time at much higher temporal resolution than currently available via conventional follow-up appointments spaced weeks apart. Furthermore, EIS holds promise for early detection of fracture nonunion, which currently can take over 6 months to diagnose, as it is more sensitive to the early stages of hematoma and cartilage than traditional techniques and can thus more quickly identify stalled healing.

In this paper, we have presented complete histologic, radiographic, and µCT analyses with our data to show that our local EIS measurements can evaluate the fracture callus with high sensitivity. Use of both external fixators and bone plates demonstrates the value of EIS in different models with different rates and progression of healing, with even the smallest of fixation implants providing a convenient platform for adding sensors and instrumentation. This establishes EIS as a technology that is amenable to miniaturization and can thus be translated and easily integrated into current fracture management strategies in the clinic. Our study lays the groundwork for instrumented implants, which could be used during surgical intervention to provide physicians with additional quantitative information during post-operative monitoring and help guide clinical care decisions.

## Materials and Methods

### Study Design

Our goal for this study was to assess the feasibility of EIS as a quantitative tool to monitor fracture healing. From previous studies, we hypothesized that EIS measurements would correlate well with biological indicators of healing, such as callus composition, X-ray radiographs, and µCT indices (i.e. bone mineral density). We conducted a controlled laboratory experiment in a murine fracture model, stabilizing fractures with an external fixator or bone plate and inserting custom sensors in the fracture gap. Euthanasia time points were selected to provide information about callus composition over the course of healing.

### Impedance Measurement System

Two-point impedance measurements were taken at the following frequencies (Hz): 20, 100, 1k, 5k, 10k, 15k, 25k, 50k, 75k, 100k, 250k, 500k, 750k, 1M. Impedance magnitude (|Z|) and phase (θ) were recorded from an Agilent E4980AL Precision LCR Meter with a 100 mV sine wave output signal, and converted to electrical resistance (R) and reactance (X). 100 mV was chosen to be below the threshold for electrolysis of water, and as high as necessary to obtain good signal-to-noise ratio. At each measurement time point, data was averaged over three sets of measurements taken for each sample, with each set involving 5 impedance measurements at the 14 frequencies listed above.

### Sensor Development

#### External Fixator Model

250 µm-diameter sensor pins were fabricated using an FR4 (a glass-reinforced epoxy laminate material) substrate with electroless nickel immersion gold (ENIG) plated electrodes (127 µm diameter) spaced 500 µm apart (PCB Universe, Vancouver, WA). Smaller sensors were then made for a subsequent study to minimize disruption of healing. 25.4 µm-diameter platinum (Pt) wires insulated with 1.27 µm isonel (A-M Systems, Sequim, WA) were used to create sensors small enough for implantation in a fracture model involving the mouse tibia, which is approximately 1 mm in diameter. Two wires were cut to roughly 50 mm in length, and recording sites were exposed by burning off short 175 µm sections of insulation in the center of each wire using a CO_2_ laser. The two wires were offset so the electrodes were spaced 0.5 mm apart, then twisted together to form a single sensor. The wires were coiled on either side of the electrodes to allow for strain relief.

#### Bone Plate Model

Flexible sensors were fabricated on 115 µm-thick polyimide (AltaFlex, Santa Clara, CA) to be affixed to the bone plate. Epo-Tek H20E silver (Ag) epoxy (Epoxy Technology, Inc., Billerica, MA) was used to conductively adhere 25.4 µm-diameter platinum (Pt) wires to Electroless Nickel Immersion Gold (ENIG)-plated pads on the polyimide sensor board. These Pt wires were attached at the center of the bone plate to serve as electrodes directly in the fracture gap. A long serpentine cable extended from the proximal end of the plate, leading to a set of connection pads that interfaced with our measurement instrumentation. The entire sensor was then coated with 6-15 µm of Parylene-C at room temperature through a chemical vapor deposition process using an SCS Labcoter^®^ 2 Parylene Deposition System (Specialty Coating Systems, Indianapolis, IN) as an insulator and for biocompatibility. A sharp razor blade was used to cut off the ends of the Pt wires to expose recording sites (700 µm^2^) spaced 0.5 mm. Medical-grade Epo-Tek 353ND epoxy was used to attach the two Pt wires and to secure the polyimide board to the proximal half of the bone plate.

Prior to implantation, fixation instrumentation was autoclaved and sensors were sterilized by immersing in 70% EtOH under ultraviolet light for at least 1 hour.

### Mouse Models

Approval was obtained from the University of California, San Francisco (UCSF) Institutional Animal Care and Use Committee (IACUC) prior to performing these mouse studies, and this report adheres to ARRIVE Guidelines for reporting animal research *(50)*. Rodent models of fracture repair have been used since 1984 *(51)* and are well-established as preclinical models that provide insight into human fracture healing *(52)*. Mice were housed in cages (≤5 mice/cage) with paper/cellulose bedding and easy access to food and water, and were monitored for signs of stress (low activity, unkempt fur, weight loss). For both models below, mice were anesthetized with an intraperitoneal (IP) injection of ketamine:dexmedetomidine (1:1) for surgery, and revived with an IP injection of atipamezole. For pain relief, buprenorphine was injected subcutaneously immediately following surgery, and subsequently at four and twenty four hours post-surgery per the IACUC protocol. Mice were given a prophylactic dose of enrofloxacin antibiotic following surgery and an additional dose on the subsequent two days.

#### External Fixator Model

Fractures were made in the mid-diaphyses of wild-type C57BL/6J mice (male, 22-29g, 12-16 weeks old) and stabilized using custom-designed external fixators. The healing timeline for this model has been previously examined in detail *(53)*. Sensors were carefully placed in the fracture gap, with skin sutured closed around the ends of the sensors, leaving wire ends available for connection to measurement instrumentation. Mice were anesthetized twice weekly using isoflurane mixed with oxygen to track healing progression with impedance measurements on days 0, 4, 7, 11, 14, 18, 21, 25, and 28. A thin, rigid acrylic sheath was placed over the external fixator to protect the sensor in between measurements. Eleven mice were implanted with 250 µm sensor pins in two different models of post-injury tissue changes – a 2 mm defect was created by an osteotomy (N=5) and a 0.5 mm defect was created by slight distraction of the proximal and distal bone ends (N=6). A 1 V sine wave output signal was used for this study. Mice were euthanized at days 7 (N=4), 14 (N=4), and 28 (N=3) for histology to reflect varying degrees of healing. Six mice were implanted with thin 56 µm Pt wire sensors in a 0.5 mm defect; 1 mouse was found dead on day 4 and taken out of the study. Mice were euthanized at day 28 for histology to allow for healing to progress to a late enough time point to diagnose proper healing or lack of healing.

#### Bone Plate Model

Ten wild-type C57BL/6J mice (male, 24-33g, 12-20 weeks old) had their right femurs stabilized with Titanium bone plates (RISystem, Davos, Switzerland), and defects (<0.25 mm wide) were made with a single cut of a Gigly saw in the mid-diaphysis. The progression of healing with this model has been previously examined in detail *(54, 55)*. Two 700 µm^2^ electrodes spaced 0.5 mm apart were centered in the fracture gap. Surgery was performed on eight additional mice, placing a bone plate and sensor but without making a fracture, to serve as controls. In these mice, the two electrodes were placed between the bone and the surrounding muscle. While infection was not a major issue, the mice had a tendency to chew or otherwise damage the connector outside the animal. To prevent this, mice were singly housed and a stretchable serpentine cable was tunneled subcutaneously up the back of each mouse, with a small incision made between the shoulder blades as an exit point. Connection pads at this location allowed for connection to measurement instrumentation. The muscle and skin was then sutured closed over the stabilized fracture. To monitor the progression of healing, mice were anesthetized with isoflurane mixed with oxygen three times each week for measurements on days 0, 2, 5, 7, 9, 12, 14, 16, 19, 21, 23, and 26. Mice were euthanized at day 12 (N=4) and day 26 (N=6 fractured, N=8 control) for histology and µCT; these two time points were chosen to reflect the transition from stage 2-3 (day 12 – mixture of cartilage and trabecular bone) and stage 3-4 (day 26 – dominated by trabecular bone, beginning to remodel). 8 mice were taken out of the study (N=1 fracture at day 12, N=1 fracture at day 26, N=6 control at day 26) because their sensors broke at the serpentine junction due to excessive movement by the animal, as seen in radiographic images and confirmed during dissection.

### Radiography

Fluoroscopy images were taken using a Hologic Fluoroscan Premier Encore 60000 C-Arm Imaging System at every measurement time point. Following euthanasia, the femur specimens were harvested and fixed overnight before being transferred to 70% ethanol for scanning. X-ray radiography was performed using a Scanco µCT 50 high-resolution specimen scanner (Scanco Medical AG, Bassersdorf, Switzerland). Specimens were imaged with X-ray energy set to 55 kVp and 109 µA. Orthogonal images were taken first with the plate intact and then again after removing the plate and screws. Orthogonal images of the bones with the plate removed were scored by five orthopaedic surgeons (single-blind assessment) using the modified Radiographic Union Scale in Tibia (RUST) fractures, with each cortex being assigned a number from 1 to 4 (1 = no callus, 2 = callus present, 3 = bridging callus, 4 = remodeled with no visible fracture line) *(49)*. The anterior cortex was excluded as the sensor typically occluded its view on an X-ray, so total values for each specimen could range from 3 to 12 by adding up the scores on the remaining three cortices. Surgeons were also asked to clinically diagnose the fractures as union, nonunion, or suspected nonunion.

Microcomputed X-ray tomography (µCT) was performed with the entire specimen scanned at an isotropic resolution of 10 µm, with an X-ray tube potential of 55 kVp and X-ray intensity of 109 µA. Integration time was set at 500 milliseconds per projection. After scanning, 3D microstructural image data was reconstructed using Scanco cone-beam reconstruction algorithm. Density calibration of the scanner was performed using a hydroxyapatite calibration phantom provided by the manufacturer. Structural indices were calculated using Scanco evaluation software (version 6.0; Scanco Medical AG). Using this software, the fracture callus was manually delineated from surrounding tissue at its cortical surface and bone morphology was assessed in the region between the central surgical screws (single-blind assessment). The callus tissue was isolated from preexisting bone by performing manual delineation to exclude the original diaphyseal cortex and its intramedullary space. The bone volume was segmented with a lower threshold of 240 grayscale units and image noise was reduced using a Gaussian filter set at a sigma and support of 0.8 and 1, respectively. Three-dimensional microstructural indices reported include bone volume (BV), total volume (TV), trabecular number, trabecular thickness, trabecular separation, and bone mineral density (BMD). Surface rendered, three-dimensional images were generated using Scanco Ray v4.0 software.

### Histology

Samples were fixed immediately after dissection in 4% paraformaldehyde (pH 7.2) overnight at 4°C, then decalcified in Cal-Ex for 48 hours at room temperature after completion of µCT scans. Samples were then processed through serial ethanol dilutions and xylene dehydration, embedded in paraffin, and serial 10 µm sections were collected and stained with Hall’s and Brunt’s Quadruple (HBQ) stain, which stains cartilage tissue blue and bone red.

### Statistical Analysis

Longitudinal impedance data was normalized to the 2^nd^ measurement time point to allow the injury to stabilize. Univariate linear regression analyses were performed to compare end-point impedance measurements to a number of µCT indices (bone volume/total volume, bone mineral density, trabecular number, trabecular thickness, and trabecular separation). Two-tailed *t*-tests determined whether regression slopes were significantly different than zero, and significance was set at p < 0.05.

## Supplementary Materials

Fig. S1 - Original histology images of external fixator model samples (Fig. 2A-B,D-E)

Fig. S2 - Frequency response over time in all bone plate model mice

Fig. S3 - Original histology images of healing and non-healing bone plate model samples (Fig. 4A,E)

Fig. S4 - Original histology images of all bone plate model samples (Fig. 5)

Fig. S5 - X-ray and µCT images of all bone plate model fracture samples

Fig. S6 - Histology image for control mouse

Fig. S7 - Goodness of fits for data fit to equivalent circuit model

Fig. S8 - Clinically-relevant frequencies of operation with significant correlation between impedance data and µCT

Table S1 - Modified RUST scores of bone plate model samples

Table S2 - Quantified µCT indices for bone plate model femur samples

Table S3 - Resulting R^2^ and p values from regression analyses comparing impedance to quantified µCT indices

## Acknowledgements

We thank Konlin Shen, Camilo Diaz-Botia, and Travis Massey for assistance with parylene-coating sensors. We thank Alfred Li for performing µCT on our specimens, which was conducted at the Bone Imaging Research Core at the San Francisco Veterans Affairs Medical Center and UCSF P30 Core Center for Musculoskeletal Biology and Medicine (CCMBM). We thank the Ella Maru Studio for creating the images in Fig. 1A and 3A. We thank Ashraf El Naga, R. Trigg McClellan, Eric Meinberg, and Zachary Working for providing clinical RUST scores for X-ray data. We thank Ralph Marcucio for reviewing data and engaging in helpful discussions, and Gina Baldoza for her work in grant and lab administration. In addition, we would like to acknowledge the Orthopaedic Trauma Institute at the Zuckerberg San Francisco General Hospital, the Berkeley Sensor and Actuator Center, the Swarm Lab at UC Berkeley, and UCSF Surgical Innovations for their support.

## Author contributions

M.C.L., M.M., S.T.H., C.S.B., and M.M.M. designed the research study, and M.C.L., D.H., and C.S.B. performed the rodent surgeries. M.C.L. collected and analyzed the data and drafted the manuscript, while M.M., S.T.H., C.S.B., and M.M.M. provided conceptual advice and critically revised the paper. M.C.L., M.M., S.T.H., C.S.B., and M.M.M. prepared supporting grants. All authors have read and approved the final submitted manuscript.

## Data and materials availability

All data needed to evaluate the conclusions are present in the paper and/or the Supplementary Materials. Additional information related to this paper may be requested from the authors.

